# A personalized anti-cancer vaccine for melanoma based on an approved vaccine against measles, mumps, and rubella

**DOI:** 10.1101/2021.06.07.447462

**Authors:** Erkko Ylösmäki, Manlio Fusciello, Arttu Uoti, Sara Feola, Beatriz Martins, Karri Aalto, Firas Hamdan, Jacopo Ciaro, Tapani Viitala, Vincenzo Cerullo

**Author notes:** Corresponding authors: Erkko Ylösmäki (E.Y.), Vincenzo Cerullo (V.C.).

## Abstract

Common vaccines for infectious diseases have been repurposed as cancer immunotherapies. Intratumoural administration of these repurposed vaccines can induce immune cell infiltration into the treated tumour. Here, we have used an approved trivalent live attenuated measles, mumps, and rubella (MMR) vaccine in our previously developed PeptiENV anti-cancer vaccine platform. Intratumoural administration of this novel MMR-containing PeptiENV anti-cancer vaccine significantly increased both intratumoural as well as systemic tumour-specific T cell responses. In addition, PeptiENV therapy, in combination with immune checkpoint inhibitor therapy, significantly improved tumour growth control and survival as well as increased the number of mice responsive to immune checkpoint inhibitor therapy.

## Introduction

The measles, mumps, and rubella (MMR) vaccine is a trivalent vaccine that contains live attenuated strains of measles, mumps, and rubella viruses. MMR vaccines are indicated for the routine immunization of children for the prevention of measles, mumps, and rubella. Recently, common vaccines for infectious diseases, such as a seasonal influenza vaccine, rotavirus vaccines and a vaccine against yellow fever, have been repurposed as intratumoural immunotherapies for the modulation of the tumour microenvironment (TME) ^1–3^. By intratumoural administration of these viral vaccines, they were able to elicit immunostimulatory effects, such as enhancement of infiltration of cytotoxic T cells (CTLs), natural killer (NK) cells and CD4^+^ Th1 T helper cells, reduction of T regulatory cells (Tregs) and in some cases, exert oncolytic properties ^1–5^. Immune checkpoint inhibitors (ICIs), a novel class of therapeutic antibodies targeting immune checkpoint molecules, such as programmed death 1 (PD-1), programmed death ligand 1 (PD-L1), and cytotoxic T lymphocyte-associated antigen 4 (CTLA-4), block the negative feedback systems within the tumour microenvironment to activate pre-existing anti-tumour immune responses ^6^. ICIs have demonstrated induction of long-term tumour regression and durable responses in some cancer patients, with response rates of 10-25% in the majority of cancers ^7^. The common feature of the patients responding to ICI therapy seems to be that they have an existing anti-tumour immunity and immune cell infiltration in the tumour tissue already prior to ICI therapy ^8, 9^. This pre-exiting anti-tumour immunity can then be enhanced and rendered functional by the ICI therapy. However, the remaining 75%-90% of the patients are not responding due to the lack of immune cell infiltration or other immune-suppressive aspects of the tumour microenvironment (TME) ^10^. As a consequence, novel therapies that can modulate the TME and attract tumour-specific CD8^+^ T cells into tumours to increase the number of patients benefiting from the ICI therapy are much needed.

Here we tested the use of an FDA/EMA approved MMR vaccine (trade name Priorix) in our recently developed peptide-based cancer vaccine platform for enveloped viruses PeptiENV ^11^ to efficiently broaden the MMR vaccine-driven immune responses to include tumour antigens. Intratumoural administration of PeptiENV Priorix significantly increased the number of tumour-infiltrating tumour-specific T cells as well as systemic tumour-specific T cells. In addition, treatment with PeptiENV Priorix suppressed tumour growth in two murine models of melanoma. This platform was then tested in combination with ICI therapy. Although PeptiENV Priorix was efficient in controlling tumour growth as a monotherapy, combining PeptiENV Priorix with an ICI against programmed death 1 (PD-1) induced significant tumour growth control and increased the number of mice responding to ICI therapy. The elegance of the PeptiENV platform is the introduction of antitumour immunity-inducing peptides non-genetically to the MMR vaccine, making this approach highly suitable for personalized immunotherapeutic approaches that rely on the identification of patient-specific neo-antigens.

## Material and methods

### Cell lines

B16F10.9/K1 cell line was kindly provided by Ludovic Martinet (Inserm, France) and was cultured in high glucose DMEM supplemented with 10% FBS, 1% L-glutamine and 1% penicillin/streptomycin. The cell line B16.OVA, a mouse melanoma cell line expressing chicken ovalbumin (OVA), was kindly provided by Prof. Richard Vile (Mayo Clinic, Rochester, MN, USA). B16.OVA cells were cultured in RPMI with 10% FBS (Life Technologies), 4 mM L-glutamine (Life Technologies), 1% penicillin/streptomycin and 5mg/mL of geneticin (Life Technologies). Murine dendritic cell line JAWSII was purchased from ATCC and was cultured in alpha minimum essential medium with 20% FBS (Life Technologies), ribonucleosides, deoxyribonucleosides, 4 mM L-glutamine (Life Technologies), 1 mM sodium pyruvate (Life Technologies), and 5 ng/ml murine GM-CSF (PeproTech, USA).

### MRR vaccine

Priorix vaccine containing the live attenuated Schwarz measles, RIT 4385 mumps (derived from Jeryl Lynn strain) and Wistar RA 27/3 rubella strains of viruses (GlaxoSmithKline Biologicals s.a.) was purchased from the University Pharmacy Ltd. (Helsinki, Finland). A single vaccination dose of Priorix contains minimum of 10^3.0^ cell culture infectious dose 50% (CCID_50_) of the Schwarz measles, 10^3.7^ CCID_50_ of the RIT 4385 mumps and 10^3.0^ CCID_50_ of the Wistar R 27/3 rubella virus strains according to the vaccine provider.

### Peptides

The following peptides were used in this study: GRKKRRQRRRPQRWEKISIINFEKL, RWEKISIINFEKL, SIINFEKL (containing an MHC class I-restricted epitope from chicken ovalbumin, OVA_257-264_). GRKKRRQRRRPQRWEKISVYDFFVWL (containing an MHC class I-restricted epitope from tyrosinase-related protein 2, Trp_2180–188_). All peptides were purchased from Zhejiang Ontores Biotechnologies (Zhejiang, China).

### PeptiENV Priorix complex formation

One vaccination dose of Priorix reconstituted in sterile water for injection was complexed with 3 nmol of CPP-extended peptides (GRKKRRQRRRPQRWEKISIINFEKL or GRKKRRQRRRPQRWEKISVYDFFVWL) or 3 nmol of control peptide (RWEKISIINFEKL).

### Surface plasmon resonance

Measurements were performed using a multi-parametric SPR Navi™ 220A instrument (Bionavis Ltd, Tampere, Finland). Phosphate buffered saline (PBS) (pH 7.4) was used as a running buffer during virus adsorption on the SPR sensor surface and during peptide-virus interaction measurements. A constant flow rate of 20 μL/min was used throughout the experiments, and temperature was set to +20°C. Laser light with a wavelength of 670 nm was used for surface plasmon excitation. A linear polyethyleneimine (LPEI, M_w_ = ~2 500 g/mol) coated Au sensor was used for the affinity measurements. LPEI renders the Au sensor slide highly positively charged which allows for efficient immobilization of measles, mumps, and rubella viruses onto the sensor. For testing the interaction between peptides and viral envelopes, 100 μM of the tested peptides (CPP-OVA or OVA control peptide without the CPP attachment moiety as a non-interacting control) were injected into a virus-coated channel and into an uncoated flow channel. The final SPR response for the peptide-virus interaction was then obtained by subtracting the SPR response in the reference channel from the measured response in the virus-coated channel.

### JAWS II dendritic cell line cross-presentation experiments

JAWSII cells were seeded in 24 well plates (2.5×10^5^ cells/well) and pulsed with PeptiENV-OVA Priorix, Priorix mixed with OVA control peptide without the CPP attachment moiety (RWEKISIINFEKL), Priorix alone or left unpulsed. After 24h, cells were collected by scraping and stained with APC-conjugated anti-mouse H-2K^b^ bound to SIINFEKL antibody (141606, BioLegend) and PerCP-Cy5.5-conjugated anti-mouse CD86 antibody (105027, BioLegend) and analysed by flow cytometry.

### Animal experiments

All animal experiments were reviewed and approved by the Experimental Animal Committee of the University of Helsinki and the Provincial Government of Southern Finland (license number ESAVI/11895/2019). Animals were kept in individually ventilated cages under standard conditions (12h light:dark, temperature- and humidity-controlled conditions) and received ad libitum access to water and food. Animals were monitored daily for symptoms related to distress and pain including hunched posture, overall activity/ability to move and roughness of the hair coat. Tumour dimensions were measured by calliper (largest tumour diameter and perpendicular tumour diameter) every second day, starting on the day tumours were first treated. All injections and tumour measurements were performed under isoflurane anaesthesia.

For the vaccination experiment using naïve mice, 8- to 9-week-old immunocompetent female C57BL/6JOlaHsd mice were subcutaneously treated on days 0, 2 and 15 with one vaccination dose of Priorix alone, one vaccination dose of Priorix together with OVA control peptide, one vaccination dose of PeptiENV-OVA Priorix, or phosphate buffered saline (PBS) as a mock-treated group. Animals were sacrificed at day 20 and spleens were harvested for immunological analysis. For the B16.OVA melanoma experiment, 8- to 9-week-old immunocompetent female C57BL/6JOlaHsd mice were injected in the right flank with 350.000 B16.OVA melanoma cells, and were treated 6-, 8- and 20-days post tumour implantation with one vaccination dose of Priorix alone, one vaccination dose of PeptiENV-OVA Priorix, peptides alone or PBS as a mock-treated group. Animals were sacrificed at day 24 and tumours, spleens and tumour-draining lymph nodes were harvested for immunological analysis. For the B16F10.9/K1 melanoma experiment, 8- to 9-week-old immuno-competent female C57BL/6JOlaHsd mice were injected in the right flank with 600.000 B16F10.9/K1 cells together with a 3:2 ratio (v:v) of Matrigel Basement Membrane Matrix High Concentration (Corning, USA), and were treated 7-, 9-, and 21-days post tumour implantation with one vaccination dose of Priorix alone, one vaccination dose of PeptiENV-Trp2 Priorix, or left untreated (Mock). Groups receiving anti-PD-1 (InVivoMab, USA, clone RMP1-14) were injected intraperitoneally three times per week with 100 μg/dose starting at day 12 post tumour implantation. Tumour growth and survival was monitored until all mice in the control group had died or sacrificed (day 35 post-tumour implantation). For the assessment of induction of systemic anti-tumour T cell responses, surviving mice were rechallenged with 1.2×10^6^ B16F10.9/K1 cells in the left flank and followed for secondary tumour development/growth.

### Enzyme-linked immunospot (ELISPOT) assays

The amount of SIINFEKL (OVA_257-264_)-specific activated, interferon-γ secreting T cells were measured by ELISPOT assay (CTL, Ohio USA) according to the manufacturer’s instructions. Briefly, 2 μg of SIINFEKL peptide was used to stimulate the antigen presenting cells. After 3 days of stimulation, plates where stained and sent to CTL-Europe GmbH for counting of the spots.

### Flow Cytometry

The following antibodies were used in the experiments: TruStain FcX™ anti-mouse CD16/32 (101320, BioLegend), FITC anti-mouse CD8 (A502-3B-E, ProImmune), Phycoerythrin (PE) anti-mouse CD3e (550353, BD Pharmingen), Peridinin-Chlorophyll-Protein (PerCP) anti-mouse CD19 (115531, BioLegend). SIINFEKL epitope-specific T cells were studied using APC-labelled H-2Kb/SIINFEKL pentamer (F093-84B-E, ProImmune). Flow cytometric analysis were performed using a BD Accuri 6C Plus (BD Biosciences) or a BD LSRFortessa™ (BD Biosciences) flow cytometer and FlowJo software v10 (BD Biosciences) was used for data analysis.

### Statistical analysis

Statistical analysis was performed using GraphPad Prism 8.0 software (GraphPad Software, USA). For data analysis, one-way or two-way ANOVA were used. All results are expressed as the mean ± SEM.

## Results

### MMR vaccine viruses can be coated with therapeutic peptides by using a cell penetrating peptide sequence as an anchor

As all three attenuated virus strains of the MMR vaccine are enveloped viruses containing a lipid bilayer derived from the host cell membrane ^12^, we hypothesized that therapeutic peptides could be attached onto the lipid bilayer of these enveloped viruses by using our recently developed peptide-based cancer vaccine platform PeptiENV ^11^ (see **Figure 1** for a schematic presentation of the platform). For the attachment moiety, we used a cell penetrating peptide (CPP) sequence derived from human immunodeficiency virus 1 (HIV-1) Tat protein. This CPP sequence was tested by surface plasmon resonance (SPR) for its efficacy at anchoring therapeutic peptides onto the envelopes of the MMR viruses. CPP moiety-containing peptide (CPP-OVA) had a robust affinity towards viral envelopes, and the affinity was specifically attributed to the CPP moiety, as peptide without the CPP moiety (OVA control peptide) had no affinity towards the viral envelopes (**Figure 2A**).

**Figure 1.**
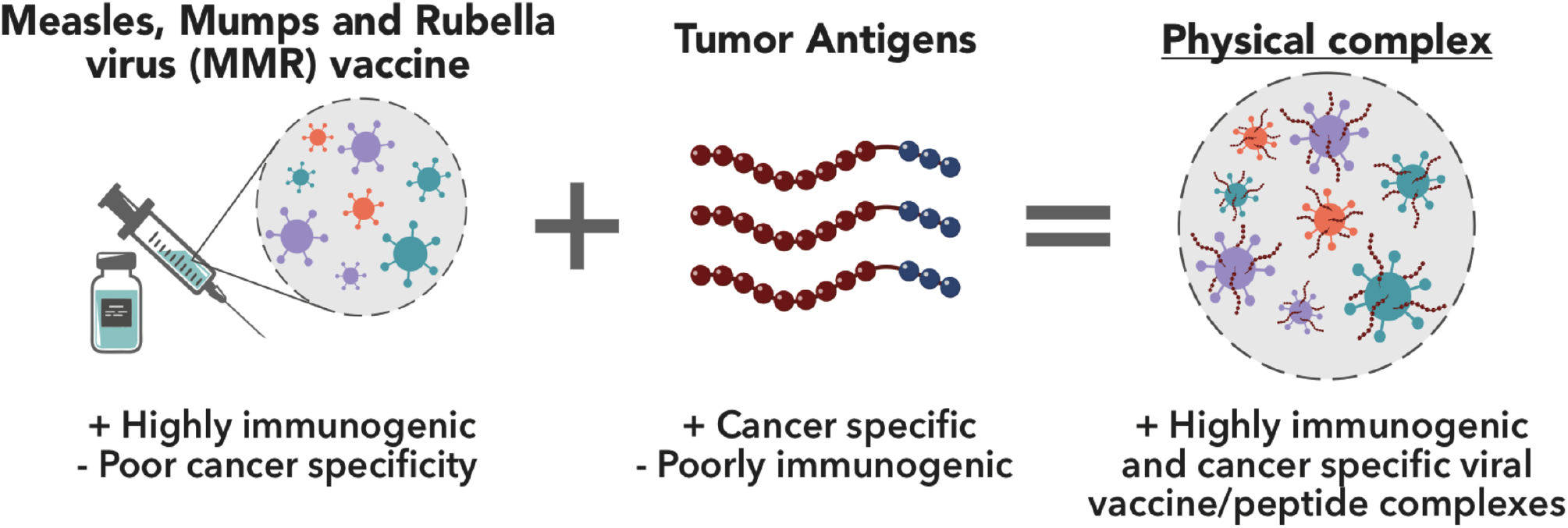
A schematic presentation of the PeptiENV cancer vaccine platform. Tumour antigens can readily be attached to the viral envelopes of measles, mumps, and rubella (MMR) vaccine viruses by using a cell penetrating peptide (CPP) sequence as an anchoring moiety. Anchor-modified peptides are complexed for 15 minutes with the MMR vaccine for allowing efficient attachment. Various different peptides, including MHC class I and II epitopes, can be delivered by the PeptiENV-platform for potent activation of antigen-presenting cells and increased antigen-specific immunological responses.

**Figure 2.**
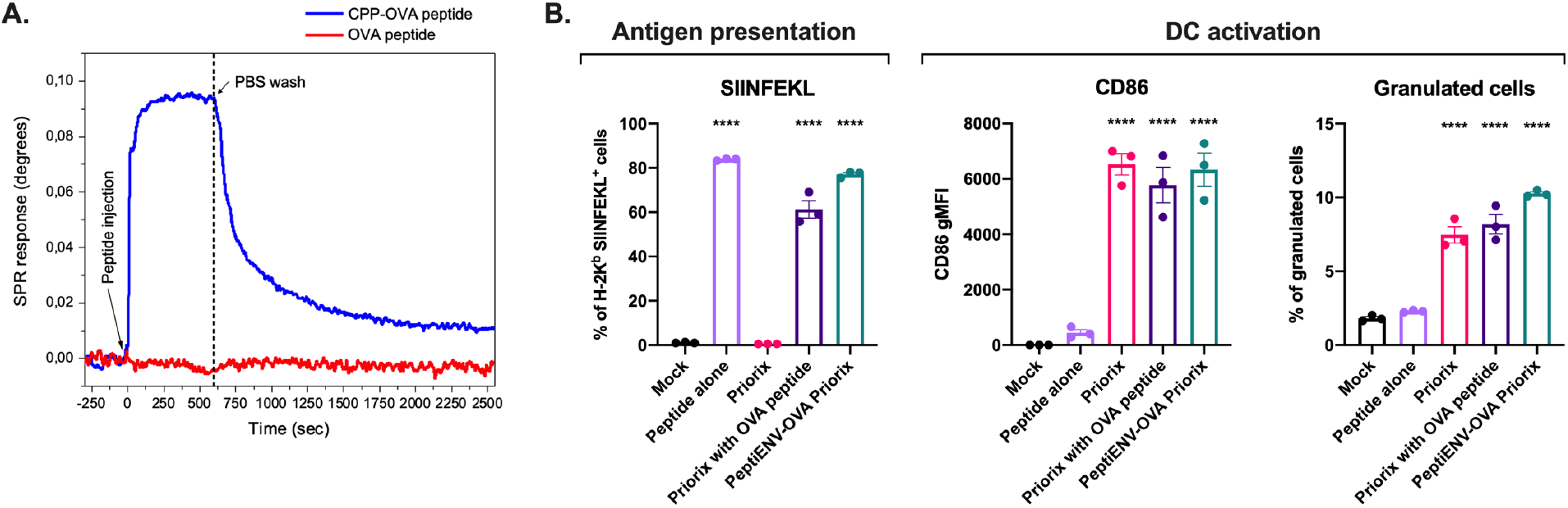
Characterization of the PeptiENV Priorix complexes. A) Surface plasmon resonance analysis of the interaction between the CPP-OVA and Priorix vaccine and OVA peptide without the attachment moiety and Priorix vaccine. B) Mouse dendritic cell line Jaws II was pulsed with PeptiENV-OVA Priorix, Priorix mixed with OVA peptide (a non-interacting control peptide), Priorix, CPP-containing SIINFEKL peptide alone or left un-pulsed (Mock). Cross-presentation was determined by flow cytometry using APC-conjugated anti-H-2Kb bound to SIINFEKL. CD86 expression and changes in morphology of the cells (as measures of dendritic cell maturation and activation) were determined by flow cytometry. Each bar is the mean ± SEM of biological triplicates. Statistical analysis was performed with one-way ANOVA. **** p< 0.0001.

### Antigen-presenting cells can readily present PeptiENV Priorix-delivered immunogenic antigens

Next, we tested whether PeptiENV platform can deliver therapeutic peptides into antigen-presenting cells (APCs) and if they can cross-present the MHC class I epitope from these peptides. Mouse dendritic cell (DC) line JAWS II was pulsed with PeptiENV-OVA Priorix, Priorix mixed with OVA control peptide, Priorix alone or CPP-OVA peptide alone, and the efficiency of cross-presentation of the mature form of the epitope (SIINFEKL) was assessed by flow cytometry (**Figure 2B**). The efficacy of SIINFEKL cross-presentation delivered by PeptiENV-OVA Priorix (77% of the cells presenting) was similar to Priorix mixed with OVA control peptide (61%) and CPP-OVA peptide alone groups (84%), indicating that peptide-coating does not affect the delivery or presentation of the peptides by the APCs. However, clear differences were seen in the ability to induce DC activation as measured by the increased expression of cluster of differentiation (CD) 86 and by the change in morphology towards more granular phenotype. Although CPP-OVA peptide was efficiently presented by the DCs, no DC activation was seen. In contrast to the peptide administration, when Priorix was administered alone, robust DC activation, but not SIINFEKL presentation was seen. When PeptiENV-OVA or Priorix mixed with OVA control peptide was administered, efficient presentation and DC activation was seen with both groups, indicating again that PeptiENV-OVA Priorix can act as a potent adjuvant for the delivered peptides.

### Vaccination with PeptiENV Priorix induces a strong immune response against the therapeutic peptide coated onto the viral envelopes of the MMR viruses

To assess the significance of physically linking the therapeutic peptides onto the envelopes of the MMR viruses, we vaccinated naïve C57BL/6JOlaHsd mice with Priorix coated with CPP-containing SIINFEKL peptide (PeptiENV-OVA Priorix) or Priorix mixed together with OVA control peptide without the CPP moiety. Mice receiving only the Priorix vaccine or PBS (mock) were used as controls. After vaccination, mice were analysed for the induction of systemic SIINFEKL-specific T cell responses by the interferon-gamma enzyme-linked immunospot (ELISPOT) assay (**Figure 3**). As expected, vaccination with Priorix or PBS did not induce any SIINFEKL-specific T cell responses. Vaccination with Priorix mixed together with SIINFEKL peptide without the CPP moiety induced SIINFEKL-specific T cell responses, but majority of the responses were moderate (average spot count 170 spots/10^6^ cells). In contrast, PeptiENV Priorix-OVA induced significantly enhanced SIINFEKL-specific T cell responses highlighting the importance of physically linking the antigen and the adjuvant (average spot count 675 spots/10^6^ cells).

**Figure 3.**
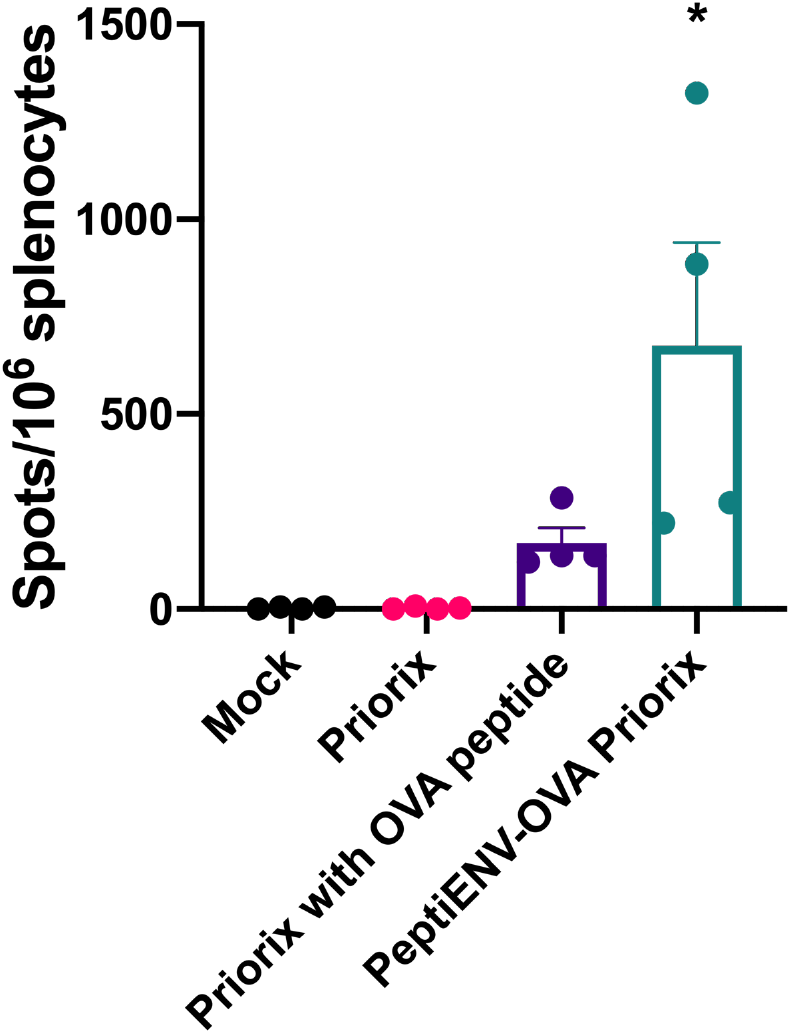
PeptiENV enhances antigen-specific T cell responses by physically linking tumour antigen and the biological adjuvant. Naive C57BL/6JOlaHsd mice (n = 4/group) were immunized with PeptiENV-OVA Priorix, Priorix mixed with OVA peptide (a non-interacting control peptide), Priorix, or PBS as a mock-treated group on days 0, 2, and 15. Five days after the last treatment, mice were sacrificed, and spleens were collected for the quantification of activated interferon-gamma (IFN-γ) secreting CD8^+^ cytotoxic T cells specific for the tumour epitope (SIINFEKL) by using a mouse interferon-gamma ELISPOT assay. Each bar is the mean ± SEM of biological quadruplicates. Statistical analysis was performed with one-way ANOVA. * p< 0.05.

### Intratumoural PeptiENV-OVA Priorix treatment induces robust systemic and intratumoural infiltration of tumour-specific T cells in a syngeneic mouse model of B16.OVA

To study the immunostimulatory potential and anti-tumour effects of the PeptiENV platform with Priorix, we used a well-established syngeneic mouse melanoma model B16 expressing chicken ovalbumin (OVA) as a model antigen ^13^. When mice bearing B16.OVA tumours were treated intratumourally with OVA-targeting PeptiENV (PeptiENV-OVA Priorix), Priorix, peptides alone or vehicle (mock), we observed only a moderate increase in tumour growth control in the PeptiENV-OVA Priorix group as compared to other treatment groups (**Figure 4A**). We set a tumour size threshold of 500 mm^3^ for defining the responders in each treatment group. Treating mice with the CPP-containing SIINFEKL peptide alone did not have an effect on tumour growth, with 2 mice defined as responders to the therapy in this group. Similarly, in Priorix- and mock-treated groups there were two responders in each group. In contrast, PeptiENV-OVA Priorix treatment had a minor effect on tumour growth with 5 mice defined as responders for the therapy. We went on to analyse whether there were any differences in immunological responses against the OVA antigen between the treatment groups, and we first assessed whether there were any differences in the infiltration of immune cells into the tumour microenvironment (TME). We did not see any differences in the number of tumour-infiltrating lymphocytes or in the number of cytotoxic CD8^+^ T cells infiltrated into the tumours between the different treatment groups. However, PeptiENV-OVA Priorix-treated mice had significantly enhanced infiltration of tumour-specific CD8^+^ T cells into the TME as compared to the tumours of Priorix-, peptide alone- or mock-treated mice (**Figure 4B**). In addition, we saw an increase of tumour-specific CD8^+^ T cells in tumour draining lymph nodes of PeptiENV-OVA-treated animals as compared to other groups (**Figure 4C**). In striking contrast to Priorix-, peptide alone- and mock-treated mice, a significant induction of a systemic OVA-specific T cell response was also seen in PeptiENV-OVA Priorix-treated mice (**Figure 4D**).

**Figure 4.**
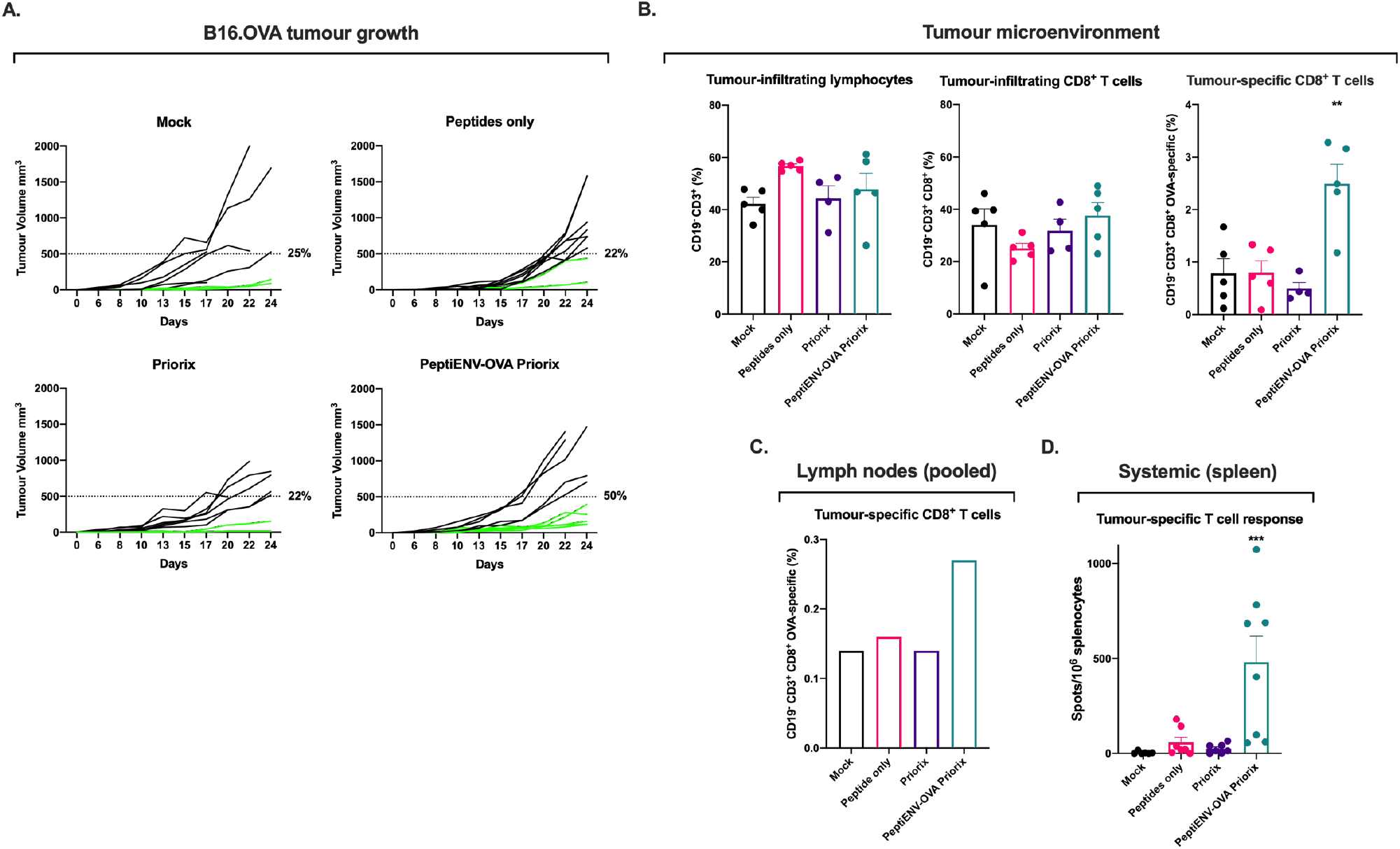
Intratumoural treatment with PeptiENV-OVA Priorix induces intratumoural and systemic tumour-specific T cell responses in a syngeneic mouse model of B16.OVA melanoma. A) PeptiENV-OVA Priorix, Priorix, peptides alone or PBS as a mock was given intratumourally 6-, 8- and 20-days post tumour implantation. Individual tumour growth curves for all treatment groups are shown. The number of mice in each group was 7–10. A threshold of 500 mm^3^ was set to define the percentage of mice responding to the different therapies (dotted line). The percentage of responders in each treatment group is shown on the right side of the dotted line. Immunological analysis of B) tumours C) lymph nodes and D) spleens of treated mice. Statistical analysis was performed with one-way ANOVA. ** p< 0.01, *** p<0.001.

### Intratumoural treatment with PeptiENV Priorix with CPP-containing Trp2 antigen increases the number of responders to anti-PD-1 therapy, greatly improves tumour control and induces systemic anti-tumour immune responses in a syngeneic mouse model of B16.F10.9/K1 melanoma

Finally, we tested the PeptiENV Priorix platform in a syngeneic mouse model of B16.F10.9/K1 melanoma using a more relevant, tumour-associated antigen from tyrosinase related protein 2 (Trp2_180–188_) in combination with anti-PD-1 immune checkpoint inhibitor therapy (ICI). B16.F10.9/K1 melanoma is a derivative of a highly metastatic B16.F10.9 melanoma with a low cell surface expression of major histocompatibility complex 1 (MHC-I) H-2Kb that was transfected with H-2Kb genes to generate H-2Kb-expressing clone K1 ^14^. The B16.F10.9/K1 clone is more responsive to cancer immunotherapies than the highly immunosuppressive parental strain B16.F10.9. Starting at 7 days post tumour engraftment, mice were treated with Priorix, anti-PD-1 alone, PeptiENV-Trp2 Priorix, Priorix in combination with anti-PD-1, PeptiENV-Trp2 Priorix in combination with anti-PD-1 or left untreated (mock). Priorix and PeptiENV-Trp2 Priorix treatments were given intratumourally, and anti-PD-1 therapy was given intraperitoneally. Here, with the more immunogenic tumour model, we set the tumour size threshold of 250 mm^3^ for defining the responders in each treatment group. In contrast to mock-treated animals, Priorix- and anti-PD-1-treated groups showed modest tumour growth control with response rates of 10% and 18%, respectively. Priorix in combination with anti-PD-1-treated animals showed moderate tumour growth control with a 45% response rate. PeptiENV-Trp2 Priorix-treated animals showed efficient tumour growth control with a response rate of 60%. Remarkably, PeptiENV-Trp2 Priorix in combination with anti-PD-1-treated animals showed the most efficient tumour growth control, with a 91% response rate; increasing the response rate for anti-PD-1 therapy from 18% to 91% (**Figure 5A**). All treatment groups increased the survival of the animals as compared to mock group (**Figure 5B**), however, treatment with PeptiENV-Trp2 Priorix- or PeptiENV-Trp2 Priorix in combination with anti-PD-1 increased most efficiently the survival of the animals, and by the time all mice in the mock group had died, 100% of the PeptiENV-Trp2 Priorix in combination with anti-PD-1-treated mice were still alive. In order to study systemic anti-tumour immunity elicited by the different treatments, surviving mice were rechallenged with a very high amount (1.2×10^6^, 2× the initial dose) of the same tumour cells into the contralateral flank. All naïve mice (n=5), used as engraftment control, showed robust tumour growth and at day 7 post rechallenge, had developed tumours with an average volume of 50mm^3^. In contrast, rechallenged mice in Priorix in combination with anti-PD-1 (n=5), PeptiENV-Trp2 Priorix (n=5) and PeptiENV-Trp2 Priorix in combination with anti-PD-1 (n=7) groups had significantly smaller tumours with average volumes of 12.6mm^3^, 17.9mm^3^ and 10.9mm^3^, respectively. In addition, 20% of rechallenged mice in Priorix in combination with anti-PD-1 and in PeptiENV-Trp2 groups did not show secondary tumour growth as compared to 43% in PeptiENV-Trp2 Priorix in combination with anti-PD-1 group, indicating an induction of a systemic anti-tumour immunity by these treatment modalities (**Figure 5C**).

**Figure 5.**
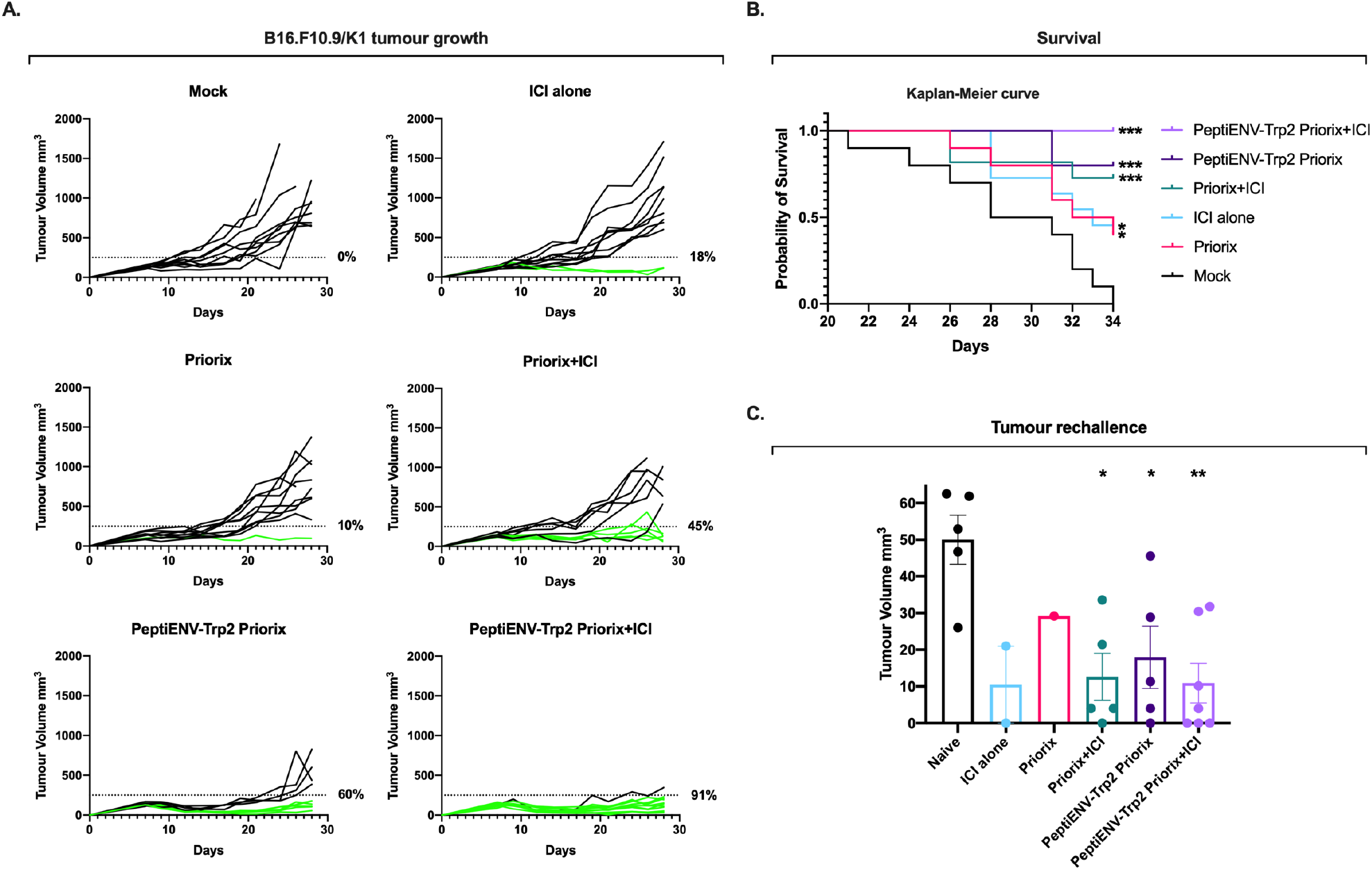
PeptiENV-Trp2 Priorix in combination with anti-PD1 improves tumour growth control and survival, and triggers a systemic anti-tumour memory response in a syngeneic mouse model of B16.F10.9/K1. A) Anti-PD-1 immune checkpoint inhibitor alone (100μg/dose given intraperitoneally three times a week, starting at day 14), Priorix alone or in combination with anti-PD-1 immune checkpoint inhibitor and PeptiENV-Trp2 alone or in combination with anti-PD-1 immune checkpoint inhibitor was given intratumourally 7-, 9-, and 21-days post tumour implantation. Individual tumour growth curves for all treatment groups are shown. A threshold of 250 mm3 was set to define the percentage of mice responding to the different therapies (dotted line). The percentage of responders in each treatment group is shown on the right side of the dotted line. The number of mice in each group was 10-11. B) Kaplan-Meier survival curve for each treatment group. C). Individual tumour volumes for rechallenged mice at day 7 after secondary tumour engraftment. Naïve mice (n=5) were used as engraftment control. Statistical analysis was performed with one-way ANOVA. * p< 0.05, ** p< 0.01, *** p<0.001.

## Discussion

In this study, we have exploited the immunostimulatory characteristics of a safe and widely used FDA and EMA-approved vaccine against measles, mumps and rubella, the MMR vaccine trade named Priorix. One vaccine dose of Priorix contains very low amounts of highly attenuated strains of measles, mumps, and rubella viruses. As all these vaccine viruses are enveloped, we set up to test the suitability of Priorix vaccine in our recently developed peptide-based cancer vaccine platform for enveloped viruses PeptiENV ^11^. By using surface plasmon resonance (SPR) analysis, we showed that cell penetrating peptide (CPP) sequence derived from HIV Tat protein can efficiently act as an attachment moiety when fused to the N-terminus of tumour antigens. Based on our earlier study, addition of the CPP sequence to the therapeutic peptides does not affect the presentation of tumour epitopes from these CPP-containing peptides ^11^. By using an immunodominant epitope from chicken ovalbumin (OVA) as a model antigen, we were able to study the characteristics of antigen presentation and concomitant dendritic cell activation by the PeptiENV platform. PeptiENV-OVA Priorix was able to efficiently deliver tumour antigens into dendritic cells and induce their robust activation. Similar characteristics was also seen with Priorix mixed with OVA control peptide lacking the CPP attachment moiety. These *in vitro* results are expected since pulsing dendritic cell monolayer in a cell culture dish does not take into account the differences in diffusion kinetics of peptides and the biological adjuvant (Priorix). Indeed, in *in vivo* vaccination setting, when the tumour antigen was physically linked to the Priorix viruses (PeptiENV-OVA Priorix) we saw significantly improved OVA antigen-specific T cell responses as compared to the situation where the peptide and Priorix did not have a physical linkage (Priorix with OVA peptide). The clear difference in OVA antigen-specific T cell responses between the non-linked (Priorix with OVA peptide) and the physically linked (PeptiENV-OVA Priorix) complexes is likely due to the limited amount of Priorix as the biological adjuvant, and in situations where the adjuvant is limited (such as when using low viral doses or intravenous administration) it might be highly beneficial to physically link the antigen to the adjuvant to increase the changes of them to reach the same antigen presenting cell for efficient induction of T cell responses.

After establishing the parameters of the physical complex and immunostimulatory potential of PeptiENV-OVA Priorix, we moved on to test the effects of the PeptiENV Priorix platform on tumour growth and OVA antigen-specific T cell induction in a murine melanoma model of B16.OVA (expressing chicken OVA as a model antigen). Although we saw only a modest tumour growth control with PeptiENV-OVA Priorix, we saw significant increase in systemic and intratumoural OVA antigen-specific T cells as well as increased number of OVA antigen-specific T cells in tumour-draining lymph nodes. These results prompted us to test the PeptiENV Priorix platform in combination with immune checkpoint inhibitors as these therapies can benefit from the increased immune cell infiltration into the TME ^9^. To test the effects of the PeptiENV Priorix platform in combination with anti-PD-1 ICI on tumour growth and tumour-specific T cell induction, we used a syngeneic mouse model of B16.F10.9/K1 melanoma together with a more relevant tumour epitope from endogenous tumour-associated antigen Trp2_(180–188)_. Interestingly, both Priorix and anti-PD-1 monotherapies had a very minimal effect on tumour growth control, but the combination therapy (Priorix + ICI) had an enhanced effect on tumour growth control. However, already as a monotherapy, PeptiENV-Trp2 Priorix had a robust effect on tumour growth control, and when combined with anti-PD-1 therapy, the antitumoural effects were further enhanced.

Vaccine strains of measles and mumps have been extensively tested for oncolytic cancer virotherapy ^15, 16^. However, the number of viral particles given intratumourally have typically been at the level of 10^6^ infectious viral particles per dose, which is 100 to 1000 times more than what is used in this study (one vaccine dose of Priorix contains approximately 10^3.0^ CCID_50_ and 10^3.7^ CCID_50_ of measles and mumps, respectively). Indeed, we wanted to use the same FDA/EMA-approved dose that has safely been used in vaccination programmes worldwide for vaccinating against measles, mumps, and rubella. With the dose used, we did not see any oncolysis-driven anti-tumoural effects by the Priorix vaccine, however, when combined with the PeptiENV platform, Priorix acted as an outstandingly potent biological adjuvant for the attached tumour antigen peptides and was able to induce anti-tumoural effects through the stimulation of tumour antigen-specific T cell immunity.

Recently, common live attenuated vaccines for various infectious diseases, have been repurposed as intratumoural immunotherapy for the modulation of the tumour microenvironment (TME) ^1, 3^. As an example, Aznar et al. successfully repurposed a live attenuated yellow fewer virus vaccine strain 17D as an intratumourally administered cancer immunotherapy ^1^. However, the dose used was much higher than the FDA/EMA-approved dose for vaccination against the yellow fever virus (intratumoural dose of 4×10^6^ plaque forming units vs. vaccination dose of 10^3^ infectious units). As the yellow fewer virus is an enveloped virus, it is intriguing to hypothesize that by using the yellow fever vaccine in the PeptiENV platform, one could further enhance the number of tumour-specific CD8^+^ T cells and thus further increase the anti-tumour efficacy of the yellow fewer virus vaccine strain 17D and possibly, allow for the reduction in the amount of vaccine virus used to minimize the risk of adverse events.

Recent studies have suggested that vaccination with the MMR vaccine may induce some cross-protective immunity against coronavirus disease 2019 (COVID-19) ^17–20^, and the efficacy of the MMR-induced cross-protection against COVID-19 is now being clinically tested (NCT04333732). However, the efficacy of the MMR vaccine to provide protection against COVID-19 could be significantly improved by the use of the PeptiENV platform with SARS-CoV-2-specific antigen peptides.

In summary, we demonstrated that an FDA/EMA-approved vaccine against measles, mumps, and rubella can be used as a potent biological adjuvant in the PeptiENV cancer vaccine platform. Treatment with PeptiENV Priorix induced robust tumour-specific T cell responses, and when combined with immune checkpoint inhibitor therapy, the number of mice responding to ICI therapy was markedly increased. As more safe, personalized cancer therapies are needed for the beneficial modulation of the tumour microenvironment for enhanced anti-tumour efficacy and for boosting of immune checkpoint inhibitor therapies, there is a rationale to convert existing FDA/EMA-approved vaccines for infectious diseases to be used in these personalized and targeted approaches. The PeptiENV Priorix vaccine platform could be rapidly taken into clinical testing, particularly in combination with immune checkpoint inhibitor therapies.

## Declarations

### Availability of data and material

All data relevant to the study are included in the article and are available upon reasonable request.

### Authors’ contributions

E.Y., M.F., A.U., and V.C. conceived and planned the experiments. E.Y., M.F., A.U., S.F., F.H., J.C., K.A., and T.V. carried out the experiments. E.Y., M.F., A.U., B.M., S.F., F.H., J.C., K.A., T.V. and V.C. contributed to the interpretation of the results. E.Y. took the lead in writing the manuscript. All authors provided critical feedback and helped shape the research, analysis, and manuscript.

